## Supplementary material for "Multi-tiered actions of *Legionella* effectors to modulate host Rab10 dynamics": Key Resources Table

| REAGENT or RESOURCE | SOURCE | IDENTIFIER |
|---|---|---|
| **Antibodies** | | |
| anti-FLAG (M2) mouse monoclonal antibody | Sigma | Cat# F1804 |
| anti-HA mouse monoclonal antibody | MBL | Cat# M132-3 |
| anti-HA rabbit monoclonal antibody | MBL | Cat# 561 |
| anti-Ub antibody (FK2) | Enzo | Cat# BML-PW8810 |
| anti-Ub antibody (P4D1) | Santa Cruz | Cat# sc-8017 |
| anti-GFP rabbit polyclonal antibody | MBL | Cat# 598 |
| anti-His monoclonal antibody | Novagen | Cat# 70796-3 |
| anti-Myc mouse monoclonal antibody | Roche | Cat# 11 667 203 001 |
| anti-RFP rabbit polyclonal antibody | MBL | Cat# PM005 |
| anti-*Legionella pneumophila* rabbit polyclonal antibody | BioAcademia | Cat# 64-100 |
| Goat anti-mouse IgG (H+L) secondary antibody, HRP | Thermo Fisher | Cat# 62-6520 |
| Goat anti-rabbit IgG (H+L) secondary antibody, HRP | Thermo Fisher | Cat# 65-6120 |
| Alexa Fluor 488 goat anti-mouse antibody | Thermo Fisher | Cat#A-11029 |
| Alexa Fluor 488 goat anti-rabbit antibody | Thermo Fisher | Cat#A-11034 |
| Rhodamine RedX goat anti-rabbit antibody | Thermo Fisher | Cat# R6349 |
| **Chemicals, Peptides, and Recombinant Proteins** | | |
| Lipofectamine2000 | Invitrogen | Cat# 11668-019 |
| Polyethylenimine (PEI) | Polysciences | Cat# 24765-2 |
| Poly-L-lysine | Sigma | Cat# P4707 |
| Paraformaldehyde (PFA) | Sigma | Cat# 441244 |
| 4,6-diamidino-2-phenylindole (DAPI) | DOJINDO | Cat# GW094 |
| ProLongTM Diamond Antifade Mountant | Thermo Fisher | Cat# P36961 |
| cOmplete$^{TM}$ protease inhibitor Cocktail (EDTA free) | Roche (Merk) | Cat# 11873580001 |
| SigmaFast Protease Inhibitor Cocktail | Sigma | Cat# S8830 |
| Phenylmethylsulfonyl fluoride (PMSF) | Nacarai | Cat# 27327-94 |
| MG132 | Calbiochem | Cat# 474791 |
| N-Ethylmaleimide (NEM) | Sigma | Cat# E3876 |
| Ni-nitrilotriacetic acid (NTA) agarose | QIAGEN | Cat# 30210 |
| FLAG M2 magnetic beads | Sigma | Cat# M8823 |
| Myc-Trap magnetic beads | chromotek | Cat# ytma |
| RFP-Trap magnetic beads | chromotek | Cat# rtma |
| Ubiquitin, human recombinant | Boston Biochem | Cat# U-100H |
| Ubiquitin K63R, human recombinant | Boston Biochem | Cat# UM-K63R |

| | | |
|---|---|---|
| Ubiquitin mutant with K63 only, human recombinant | Boston Biochem | Cat# UM-K630 |
| UBE1, human recombinant | Boston Biochem | Cat# E-305 |
| Ubc (E2) Enzyme Kit | Boston Biochem | Cat# K-980B |
| **Critical Commercial Assays** | | |
| Silver Stain MS Kit | FUJIFILM Wako | Cat# 299-58901 |
| QuickChange II site-directed mutagenesis kit | Agilent | Cat# 200523 |
| Gibson assembly kit | New England Biolabs | Cat# E2611 |
| EndoFree Plasmid MAXI prep kits | QIAGEN | Cat# 12362 |
| **Media and serum** | | |
| N-(2-acetamido)-2-aminoethanesulfonic acid (ACES) | Sigma | Cat# 7365-82-4 |
| Minimum Essential Medium α (MEMα) | Gibco | Cat# 12571-063 |
| Dulbecco's Modified Eagle Medium (DMEM) | Gibco | Cat# 11885-084 |
| Fetal Bovine Serum (FBS) | Sigma | Cat# 172012 |
| Goat Serum | Gibco | Cat# 16210-064 |
| **Experimental Models: Cell Lines** | | |
| HeLa-FcγRII | (Arasaki et al., 2017) | N/A |
| HEK293T-FcγRII | (Arasaki and Roy, 2010) | N/A |
| **Bacterial strains** | | |
| *Legionella pneumophila* Philadelphia-1 (Lp01) | (Berger and Isberg, 1993) | NC_002942.5 |
| Lp01Δ*icmV* Δ*dotA* (Δ*dotA*) | (Zuckman et al., 1999) | N/A |
| Lp01 Δ*sidC*Δ*sdcA* | This study | N/A |
| Lp01 Δ*sidC*Δ*sdcA*Δ*sdcB* | This study | N/A |
| Lp01 Δ*sidE*Δ*sdeA*Δ*sdeB*Δ*sdeC* (Δ*sidEs*) | This study | N/A |
| Lp01 Δ*dupA*Δ*dupB* | This study | N/A |
| Lp01 Δ*dupA*Δ*sidJ*Δ*dupB*Δ*sdjA* | This study | N/A |
| Lp01 ΔΔ*lpg2149* | This study | N/A |
| Lp01 Δ*mavC*Δ*mvcA* | This study | N/A |
| *Escherichia coli* DH5α | TOYOBO | Cat# DNA-903 |
| *Escherichia coli* DH5αλ*pir* | (Zuckman et al., 1999) | N/A |
| *Escherichia coli* BL21(DE3) | NOVAGEN-MERK | Cat# 69450 |
