## Supplementary material for "Multi-tiered actions of *Legionella* effectors to modulate host Rab10 dynamics": Table S1

### Appendix 1-table 1. Plasmids used in this study.

| Plasmid ID | Plasmids | Genotype | Purpose | Reference or source |
|---|---|---|---|---|
| NA | pUC18 | Cloning vector, $Amp^r$ | Cloning | TaKaRa |
| pNH2802 #2 | pUC18-*sdeA* | Cloning vector encoding SdeA, $Amp^r$ | Cloning | This study |
| pNH2804 #2 | pUC18-*sdeA*$_{EE/AA}$ | Cloning vector encoding SdeA catalytic E860A E862A mutant, $Amp^r$ | Cloning | This study |
| NA | pHA-Ub | pHA encoding Ub, $Amp^r$ | Transfection | Gift from Michinaga Ogawa |
| pNH2175 #2 | pHA-Ub$_{AA}$ | pHA encoding Ub G75A G76A mutant, $Amp^r$ | Transfection | This study |
| pNH2156 #1 | pHA-Ub$_{Q40E}$ | pHA encoding Ub Q40E mutant, $Amp^r$ | Transfection | This study |
| pNH2276 #1 | pHA-Ub$_{Q31E}$ | pHA encoding Ub Q31E mutant, $Amp^r$ | Transfection | This study |
| pNH2277 #1 | pHA-Ub$_{Q41E}$ | pHA encoding Ub Q41E mutant, $Amp^r$ | Transfection | This study |
| pNH2288 #1 | pHA-Ub$_{Q31E\ Q41E}$ | pHA encoding Ub Q31E Q41E mutant, $Amp^r$ | Transfection | This study |
| NA | p3xHA-Ub | p3xHA encoding Ub, $Amp^r$ | Transfection | Gift from Jiazhang Qiu |
| NA | p3xHA-Ub$_{AA}$ | p3xHA encoding Ub G75A G76A mutant, $Amp^r$ | Transfection | Gift from Jiazhang Qiu |
| NA | pHA-Ub$_{K6}$ | pHA encoding Ub with Lys6 (without any other Lys residues), $Amp^r$ | Cloning | Addgene #22900 |
| pNH2177 #1 | pHA-Ub$_{No\ K}$ | pRK5-HA encoding Ub with no Lys residues, $Amp^r$ | Transfection | This study |
| NA | pEGFP-C2 | GFP-tagged protein expression vector, $Kan^r$ | Transfection | BD |
| NA | pmGFP | pEGFP-C2 encoding monomeric GFP (GFP with L222K mutation), $Kan^r$ | Transfection | (Kitao et al., 2020) |
| pNH2123 #1 | pmGFP-*mavC* | pmGFP encoding MavC, $Kan^r$ | Transfection | This study |
| pNH2140 #1 | pmGFP-*mavC*$_{C74A}$ | pmGFP encoding MavC C74A mutant, $Kan^r$ | Transfection | This study |
| NA | pET15b | hexahistidine-tagged protein expression vector, $Amp^r$ | Protein purification | Novagen |
| pNH1989 #9 | pET15b-His-*sdeA* | pET15b encoding hexahistidine-tagged SdeA, $Amp^r$ | Cloning | This study |
| NA | pET15b-His-*sdeA*$_{\Delta DUB}$ | pET15b encoding hexahistidine-tagged SdeA Δ1-199aa , $Amp^r$ | Cloning | This study |
| pNH2000 #6 | pmGFP-*sdeA* | pmGFP encoding SdeA, $Kan^r$ | Cloning | This study |

| | | | | |
|---|---|---|---|---|
| NA | pmGFP-*sdeA* $_{\Delta DUB}$ | pmGFP encoding SdeA Δ1-199aa , $Kan^r$ | Transfection | This study |
| NA | P3xFLAG-cDNA4T/O- | 3xFLAG-tagged protein expression vector, $Amp^r$ | Transfection | (Ingmundson et al., 2007) |
| NA | p3xFLAG-Rab10 | p3xFLAG encoding human Rab10A, $Amp^r$ | Transfection | This study |
| pNH2515 #1 | p3xFLAG-Rab10QL | p3xFLAG encoding human Rab10A Q68L, $Amp^r$ | Transfection | This study |
| pNH2516 #1 | p3xFLAG-Rab10TN | p3xFLAG encoding human Rab10A T23N, $Amp^r$ | Transfection | This study |
| pNH2518 #2 | p3xFLAG-Rab10KKK | p3xFLAG encoding human Rab10A K102A K136A K154A, $Amp^r$ | Transfection | This study |
| NA | p3xFLAG-CMV-10 | 3xFLAG-tagged protein expression vector, $Amp^r$ | Transfection | Sigma |
| pNH1807 #1 | p3xFLAG-*sidC* | p3xFLAG encoding SidC, $Amp^r$ | Transfection | This study |
| pNH1808 #26 | p3xFLAG- *sidC* $_{C46A}$ | p3xFLAG encoding SidC C46A mutant, $Amp^r$ | Transfection | This study |
| pNH2172 #1 | p3xFLAG-*sdcA* | p3xFLAG encoding SdcA, $Amp^r$ | Transfection | This study |
| pNH2173 #2 | p3xFLAG- *sdcA* $_{C44A}$ | p3xFLAG encoding SdcA C44A mutant, $Amp^r$ | Transfection | This study |
| pNH2109 #4 | p3xFLAG-*sdcB* | p3xFLAG encoding SdcB mutant, $Amp^r$ | Transfection | This study |
| pNH2126 #6 | p3xFLAG- *sdcB* $_{C57A}$ | p3xFLAG encoding SdcB C57A mutant, $Amp^r$ | Transfection | This study |
| pNH2286 #1 | p3xFLAG- *sdcB* $_{K518R}$ | p3xFLAG encoding SdcB K518R mutant, $Amp^r$ | Transfection | This study |
| pNH2287 #1 | p3xFLAG- *sdcB* $_{K891R}$ | p3xFLAG encoding SdcB K891R mutant, $Amp^r$ | Transfection | This study |
| pNH2289 #3 | p3xFLAG- *sdcB* $_{K518R\ K891R}$ | p3xFLAG encoding SdcB K518R K891R mutant, $Amp^r$ | Transfection | This study |
| NA | pCMV-HA-N | N-terminal HA-tagged protein expression vector, $Amp^r$ | Transfection | Clontech |
| pNH2272 # | pHA-*mavC* | pHA encoding MavC, $Amp^r$ | Transfection | This study |
| pNH2273 #1 | pHA-*mavC* $_{C74A}$ | pHA encoding MavC C74A mutant, $Amp^r$ | Transfection | This study |
| NA | pmRFP-C1 | N-terminal RFP-tagged protein expression vector, $Kan^r$ | Transfection | (Murata et al., 2006) |
| NA | pmRFP-Rab10 | pRFP encoding Rab10, $Kan^r$ | Transfection | This study |
| pNH1804 #1 | pET15b-His-*sidC* | pET15b encoding hexahistidine-tagged SidC, $Amp^r$ | Protein purification | This study |
| pNH2120 #6 | pET15b-His-*sdcB* | pET15b encoding hexahistidine-tagged SdcB, $Amp^r$ | Protein purification | This study |
| pNH2121 #9 | pET15b-His-*sdcB* $_{C57A}$ | pET15b encoding hexahistidine-tagged SdcB C57A mutant, $Amp^r$ | Protein purification | This study |
| pNH2142 #1 | pET15b-His-*mavC* | pET15b encoding hexahistidine-tagged MavC, $Amp^r$ | Protein purification | This study |
| pNH2143 #5 | pET15b-His-*mavC* $_{C74A}$ | pET15b encoding hexahistidine-tagged MavC C74A mutant, $Amp^r$ | Protein purification | This study |

| | | | | |
|---|---|---|---|---|
| pNH2144 #3 | pET15b-His-*mvcA* | pET15b encoding hexahistidine-tagged MvcA, $Amp^r$ | Protein purification | This study |
| NA | pSR47S | oriR6K, *oriT* RP4, *Kan$^r$*, *sacB* | Gene deletion | (Merriam et al., 1997) |
| pNH1990 #1 | pSR47S-Δ*sidE* | pSR47S carrying 30bp upstream and downstream regions of *sidE,* $Kan^r$ | Gene deletion | This study |
| pNH1993 #1 | pSR47S-Δ*sdeA*Δ*sdeB* | pSR47S carrying 300bp upstream and downstream regions of *sdeA-sdeB,* $Kan^r$ | Gene deletion | This study |
| pNH1994 #2 | pSR47S-Δ*sdeC* | pSR47S carrying 300bp upstream and downstream regions of *sdeC,* $Kan^r$ | Gene deletion | This study |
| pNH2213 #1 | pSR47S-Δ*dupA* | pSR47S carrying 300bp upstream and downstream regions of *dupA,* $Kan^r$ | Gene deletion | This study |
| pNH2196 #1 | pSR47S-Δ*dupB* | pSR47S carrying 300bp upstream and downstream regions of *dupB,* $Kan^r$ | Gene deletion | This study |
| pNH2214 #1 | pSR47S-Δ*dupA*Δ*sidJ* | pSR47S carrying 300bp upstream and downstream regions of *dupA-sidJ,* $Kan^r$ | Gene deletion | This study |
| pNH2197 #1 | pSR47S-Δ*dupB*Δ*sdjA* | pSR47S carrying 300bp upstream and downstream regions of *dupB-sdj,* $Kan^r$ *A* | Gene deletion | This study |
| pNH1803 #1 | pSR47S-Δ*sidC*Δ*sdcA* | pSR47S carrying 300bp upstream and downstream regions of *sidC-sdcA,* $Kan^r$ | Gene deletion | This study |
| pNH2111 #2 | pSR47S-Δ*sdcB* | pSR47S carrying 300bp upstream and downstream regions of *sdc,* $Kan^r$ *B* | Gene deletion | This study |
| pNH2139 #1 | pSR47S-Δ*lpg2149* | pSR47S carrying 300bp upstream and downstream regions of *sdcB*, $Kan^r$ | Gene deletion | This study |
| pNH2136 #3 | pSR47S-Δ*mavC*Δ*mvcA* | pSR47S carrying 300bp upstream and downstream regions of *mavC-mvcA,* $Kan^r$ | Gene deletion | This study |
| NA | pMMB207NT | Cloning vector with P*icmR* derived from RSF1010 (*oriR*), $Cm^r$ | Expression vector for *Legionella* | (Coers et al., 2000) |

| | | | | |
|---|---|---|---|---|
| pNH1884 #10 | pMMB207NT-3xFLAG | pMMB207NT to express 3xFLAG-tagged protein, $Cm^r$ | Expression vector for *Legionella* | (Kubori et al., 2018) |
| pNH2117 #1 | pMMB207-3xFLAG-*sdcB* | pMMB207NT encoding 3xFLAG-tagged SdcB, $Cm^r$ | Expression in *Legionella* | This study |
| pNH2118 #3 | pMMB207-3xFLAG-*sdcB*$_{C57A}$ | pMMB207NT encoding 3xFLAG-tagged SdcB C57A mutant, $Cm^r$ | Expression in *Legionella* | This study |
| NA | pMMB207NT-3xMyc | pMMB207NT to express 3xMyc-tagged protein, $Cm^r$ | Expression vector for *Legionella* | (Kubori et al., 2022) |
| NA | pMMB207NT-3xMyc-*sdeA* | pMMB207NT to express 3xMyc-tagged SdeA, $Cm^r$ | Expression in *Legionella* | This study |
| NA | pMMB207NT-3xMyc-*sdeA*$_{EE/AA}$ | pMMB207NT to express 3xMyc-tagged SdeA E860A E862A mutant, $Cm^r$ | Expression in *Legionella* | This study |
| pNH2236 #3 | pMMB207NT-3xMyc- *sdcB* | pMMB207NT to express 3xMyc-tagged SdcB, $Cm^r$ | Expression in *Legionella* | This study |
| pNH2237 #1 | pMMB207NT-3xMyc-*sdcB*$_{C57A}$ | pMMB207NT to express 3xMyc-tagged SdcB C57A mutant, $Cm^r$ | Expression in *Legionella* | This study |
| pNH2511 #14 | pMMB207NT-3xMyc-*sdcB*$_{K518R\ K891R}$ | pMMB207NT to express 3xMyc-tagged SdcB K518R K891R mutant, $Cm^r$ | Expression in *Legionella* | This study |
