## Supplementary material for "Multi-tiered actions of *Legionella* effectors to modulate host Rab10 dynamics": Table S2

**Appendix 2-table 2. List of primers used in this study.**

| **Primers** | | |
|---|---|---|
| **ID** | **Name** | **Sequence (5' to 3')** |
| 2100 | sdeA f (BamHI) | GCGGATCCAGATGCCTAAGTATGTCGAAGGG |
| 2101 | sdeA r (XbaI) | GCTCTAGATTAAAATCCTATAGTTTTTTTATTGG |
| 2104 | sdeA E860A E862A f | GCGAAGGCACCGCAAGTGCATTCTCCG |
| 2105 | sdeA E860A E862A r | CGGAGAATGCACTTGCGGTGCCTTCGC |
| 2548 | Ub GG75-76AA f | GTTGAGACTTCGTGCTGCTTAACTCGAGCATG |
| 2549 | Ub GG75-76AA r | CATGCTCGAGTTAAGCAGCACGAAGTCTCAAC |
| 2329 | Ub Q40E f | CCTCCTGATGAGCAGAGACTGATC |
| 2330 | Ub Q40E r | GTCTCTGCTCATCAGGAGGTATTC |
| 2935 | Ub Q31E f | GGCCAAGATCGAGGATAAGGAAGG |
| 2936 | Ub Q31E r | CCTTCCTTATCCTCGATCTTGGCC |
| 2937 | Ub Q41E f | CCTCCTGATCAGGAGAGACTGATC |
| 2938 | Ub Q41E r | GATCAGTCTCTCCTGATCAGGAGG |
| 2576 | pRK5-HA-Ub K6R f | GATCTTCGTCAGAACGTTAACC |
| 2577 | pRK5-HA-Ub K6R r | GGTTAACGTTCTGACGAAGATC |
| NA | Rab10 f (SmaI) | TCCCCCGGGCATGGCGAAGAAGACGTACGA |
| NA | Rab10 r (SmaI) | TCCCCCGGGTCAGCAGCATTTGCTCTTCC |
| 3015 | Rab10 Q68L f | GGGATACAGCAGGCCTGGAGCGATTTCAC |
| 3016 | Rab10 Q68L r | GTGAAATCGCTCCAGGCCTGCTGTATCCC |
| 3017 | Rab10 T23N f | GGAGTGGGGAAGAACTGCGTCCTTTTTCG |
| 3018 | Rab10 T23N r | CGAAAAAGGACGCAGTTCTTCCCCACTCC |
| 3007 | Rab10 K102A f | GAAAACATCAGCGCATGGCTTAGAAAC |
| 3008 | Rab10 K102A r | GTTTCTAAGCCATGCGCTGATGTTTTC |
| 3009 | Rab10 K136A f | CCTAAAGGAGCAGGAGAACAGATTGC |
| 3010 | Rab10 K136A r | GCAATCTGTTCTCCTGCTCCTTTAGG |
| 3011 | Rab10 K154A f | CTAGTGCAGCAGCAAATATAAACATC |
| 3012 | Rab10 K154A r | GATGTTTATATTTGCTGCTGCACTAG |
| 2217 | mavC f (XhoI) | GCCTCGAGCATGACAACTTCCAAGCTTG |
| 2218 | mavC r (PstI) | GCGCTGCAGTCACTTATCACGAAGAACTAACC |
| 2259 | mavC C74A f | CCACAAACAGCGCCGGAAAAGCG |
| 2260 | mavC C74A r | CGCTTTTCCGGCGCTGTTTGTGG |
| 1848 | sdeA f (XhoI) | GCCTCGAGATGCCTAAGTATGTCGAAGGG |
| 1849 | sdeA r (BamHI) | GCGGATCCTTAAAATCCTATAGTTTTTTTATTGG |
| 2712 | pET15b_sdeAΔDUB_F | GCCGCGCGGCAGCCATCCTACACTGGTTCTTCCTGGA |
| 2649 | pET15b_GA1 | ATGGCTGCCGCGCGGCAC |
| 1840 | sdeA f (PstI) | CGCTGCAGTATGCCTAAGTATGTCGAAGGG |

| | | |
|---|---|---|
| 2714 | pmGFP-sdeAΔDUB_F | TCCGGCCGGACTTGTACCTACACTGGTTCTTCCTGGA |
| 2715 | pmGFP-C2 GA1 | CCGGCCGGACTTGTACAGCTCGTC |
| 2574 | sidC f (KpnI) | CCGGGTACCAATGGTGATAAACATGGTTGACG |
| 1397 | sidC r (BamHI) | GAGGATCCCTATTTCTTTATAATTCCCGTG |
| 1408 | sidC C46A f | GATAATACCGCCCAAACAGCAGTTG |
| 1409 | sidC C46A r | CAACTGCTGTTTGGGCGGTATTATC |
| 2575 | sdcA f (KpnI) | CCGGGTACCAATGAACATGGTTGACAAAATAAAATTC |
| 2222 | sdcA r (BamHI) | GCGGATCCCTATATTGTATTCCTAACAG |
| 2579 | sdcA C44A f | GATAATACCGCTGAAACAACAGGTGAGTTATTAACC |
| 2580 | sdcA C44A r | CTCACCTGTTGTTTCAGCGGTATTATCCAGTCCTATTTC |
| 2186 | sdcB f (EcoRI) | CGGAATTCATTGAAAGACCAATTAGCC |
| 2165 | sdcB r (BamHI) | GCGGATCCCTAAGCCAGTTTATTGGATATTTC |
| 2743 | sdcB f (BamHI) | GCGGATCCTCTTGAAAGACCAATTAGCC |
| 2744 | sdcB r (SalI) | CGCGTCGACCTAAGCCAGTTTATTGGATATTTC |
| 2200 | sdcB C57A f | GGCGATAATACGGCCAAATCGGATTTAG |
| 2201 | sdcB C57A r | CTAAATCCGATTTGGCCGTATTATCGCC |
| 2951 | sdcB K518R f | GCTGAAGCTGTGGCGTCAAGAGTTCGCTACCTG |
| 2952 | sdcB K518R r | CAGGTAGCGAACTCTTGACGCCACAGCTTCAGC |
| 2953 | sdcB K891R f | GTTTTCTTTTCTGGCAGAGAAAATATAAAAACAGAC |
| 2954 | sdcB K891R r | GTCTGTTTTTATATTTTCTCTGCCAGAAAAGAAAAC |
| 2919 | mavC f (SalI) | CGGTCGACCATGACAACTTCCAAGCTTG |
| 2306 | mavC r (XhoI) | GCCTCGAGTCACTTATCACGAAGAACTAACC |
| 1396 | sidC f (XhoI) | GCCTCGAGGTGATAAACATGGTTGACG |
| 1397 | sidC r (BamHI) | GAGGATCCCTATTTCTTTATAATTCCCGTG |
| 2207 | sdcB f (NdeI) | GCTGCATATGTTGAAAGACCAATTAGCC |
| 2278 | mavC f (XhoI) | GCCTCGAGATGACAACTTCCAAGCTTG |
| 2279 | mavC r (BamHI) | GCGGATCCTCACTTATCACGAAGAACTAACC |
| 2280 | mvcA f (XhoI) | GCCTCGAGATGACAAAAATAAAACTGGAATCG |
| 2281 | mvcA r (BamHI) | GCGGATCCTTAGCTTGGCCCCTTTTTATACC |
| 1850 | sidE d1 (SacI) | GCGAGCTCCTAATTCAAACAGGACTTAC |
| 1851 | sidE d2 | GGAAGATAAAGAATTACATTACACAACTCCTG |
| 1852 | sidE d3 | TGTGTAATGTAATTCTTTATCTTCCACGG |
| 1853 | sidE d4 (XbaI) | GCTCTAGACTTTGAGAACATCACTTCCTG |
| 1858 | sdeB d1 (SacI) | GCGAGCTCCCGGAATCCGAACGTAC |
| 1862 | sdeBA d2 | TAAGGTCAATTACATTTTACTTTCTCCCAAGCG |
| 1863 | sdeBA d3 | GGGAGAAAGTAAAATGTAATTGACCTTAACCC |
| 1857 | sdeA d4 (XbaI) | GCTCTAGAGAGGTAATTTCATCTCAAAG |
| 1983 | sdeC d1 (SacI) | GCGAGCTCCACACCAATCGGAAGGTTTC |
| 1984 | sdeC d2 | AAAAAATACTCCTTACATTTTACTTTCTCCCAAAC |

| | | |
|---|---|---|
| 1985 | sdeC d3 | GGGAGAAAGTAAAATGTAAGGAGTATTTTTAAGTG |
| 1986 | sdeC d4 (XbaI) | GCTCTAGAGATTGTATTCCCATTCCGG |
| 2963 | dupA d1 (SacI) | GCGAGCTCGTTTCTCATCAAGCTGCTGC |
| 2964 | dupA d2 | GCCTCATAACCTCTTCACATGGTATGAGCCAAACC |
| 2965 | dupA d3 | GGTTTGGCTCATACCATGTGAAGAGGTTATGAGGC |
| 2696 | dupA d4 (XbaI) | GCTCTAGAGGTTGTGCTTGTAGTCG |
| 2684 | dupB d1 (SacI) | GCGAGCTCCTGATGTGTTGGAAGTGGCGG |
| 2685 | dupB d2 | ACTTTAAATAAAACAGGCTACATCGTATGAGCTAACCC |
| 2686 | dupB d3 | GGGTTAGCTCATACGATGTAGCCTGTTTTATTTAAAG |
| 2687 | dupB d4 (XbaI) | GCTCTAGACAACTCAAGTCTGGTATCCTC |
| 2697 | dupA sidJ d2 | ATTCGTTTTATCACATGGTATGAGCCAAACC |
| 2698 | dupA sidJ d3 | GGTTTGGCTCATACCATGTGATAAAACGAATACCC |
| 2699 | sidJ d4 (XbaI) | GCTCTAGACTTTCTCCCAAGCGAATATTTG |
| 2688 | dupB sdjA d2 | ACAATTGAAATGACCCTTTCACATCGTATGAGCTAACCC |
| 2689 | dupB sdjA d3 | GGGTTAGCTCATACGATGTGAAAGGGTCATTTCAATTG |
| 2690 | sdjA d4 (XbaI) | GCTCTAGACAATAAGATATTATTGACCG |
| 1384 | sdcA d1 (XbaI) | GCTCTAGAATAATAGGCACAATGGTCTCC |
| 1385 | sdcAsidC d2 | GATTTCACTCTTACCTACATCACCCTATGC |
| 1386 | sdcAsidC d3 | AGGGTGATGTAGGTAAGAGTGAAATCACTG |
| 1387 | sidC d4 (XbaI) | GCTCTAGAGAGGACGTTTGGGCTGAGGAG |
| 2196 | sdcB d1 (SacI) | GCGAGCTCGTATGGGTATAGCATCAGAG |
| 2197 | sdcB d2 | ATCCAGCTACAATTTCGTTATACTCTTTC |
| 2198 | sdcB d3 | GTATAACGAAATTGTAGCTGGATAGTTTAACCG |
| 2199 | sdcB d4 (XbaI) | GCTCTAGAGGATTGTCTTTGGACAGGATG |
| 2272 | lpg2149 d1 (SacI) | GCGAGCTCCTGAAATTAACCCTGAATTGGC |
| 2273 | lpg2149 d2 | CCTCATTTCAATTACATAAATCAAACTCCTTCG |
| 2274 | lpg2149 d3 | TTGATTTATGTAATTGAAATGAGGATGGGG |
| 2275 | lpg2149 d4 (XbaI) | GCTCTAGACGCTGTCAAATCTTGTAGG |
| 2263 | mavC d1 (SacI) | GCGAGCTCCAACACAGGGTATGAATAGC |
| 2264 | mavC mvcA d2 | GATTTTTATTGTTTTATTGATTACATATTAACCTCACTGATAG |
| 2265 | mavC mvcA d3 | CAGTGAGGTTAATATGTAATCAATAAAACAATAAAAATC |
| 2266 | mvcA d4 (XbaI) | GCTCTAGAGTCCATGGCATCAAGCGC |
| 2204 | sdcB f (BamHI) | GCGGATCCTCTTGAAAGACCAATTAGCC |
| 2205 | sdcB r (XbaI) | GCTCTAGACTAAGCCAGTTTATTGGATATTTC |
| 2341 | pMMB207-PicmR_2 | AAGCTTGGCTGTTTTGGCGGATG |
| 2681 | pMMB207-PicmR-3xmyc_GA1 | ACCCAGATCTTCTTCAGAGATGAGCTTCTGTTCACCTAA |
| 2658 | pMMB207-3xMyc-SdeA_F | TGAAGAAGATCTGGGTATGCCTAAGTATGTCGAAGGG |
| 2659 | pMMB207-3xMyc-SdeA_R | CAAAACAGCCAAGCTTTTAAAATCCTATAGTTTTTTATTGGATTC |
